# An alternative STAT signaling pathway acts in antiviral immunity in *Caenorhabditis elegans*

**DOI:** 10.1101/110940

**Authors:** Mélanie Tanguy, Louise Véron, Przemyslaw Stempor, Julie Ahringer, Peter Sarkies, Eric A. Miska

**Author notes:** co-corresponding author. (EAM) (PS).

## Abstract

Across metazoans, innate immunity is vital in defending organisms against viral infection. In mammals, antiviral innate immunity is orchestrated by interferon signaling, activating the STAT transcription factors downstream of the JAK kinases to induce expression of antiviral effector genes. In the nematode *C. elegans*, which lacks the interferon system, the major antiviral response so far described is RNA interference but whether additional gene expression responses are employed is not known. Here we show that, despite the absence of both interferon and JAK, the *C. elegans* STAT homologue STA-1 orchestrates antiviral immunity. Intriguingly, mutants lacking STA-1 show increased resistance to antiviral infection. Using gene expression analysis and chromatin immunoprecipitation we show that, in contrast to the mammalian pathway, STA-1 acts as a transcriptional repressor. Thus STA-1 might act to suppress a constitutive antiviral response in the absence of infection. Using a reverse genetic screen we identify the SID-3 as a kinase upstream of STA-1 in the response to infection. Together, our work identifies a novel STAT regulatory cascade controlling its activity in antiviral resistance, illustrating the complex evolutionary trajectory displayed by innate immune signaling pathways across metazoan organisms.

## Introduction

RNA viruses, a highly diverse family known to infect organisms of almost all kingdoms of life, also represent an important burden on human health. Indeed, their highly mutagenic and adaptive nature is an ever-growing challenge for diagnostic and treatments and the underlying understanding of host-pathogen interaction. The first and critical step in mounting successful antiviral defence is the conserved innate immune response, however its complexity is yet to be fully apprehended. Several pathways have been described in various organisms that differ in their presence or relative importance. To date, the main pathways can be divided in two categories: protein based innate immunity and RNA based innate immunity. Jawed vertebrates rely mostly on the powerful interferon (IFN) system, whereas most other eukaryotes take advantage of the RNA interference (RNAi) machinery.

In mammals, the initiation of the interferon response in response to RNA viruses depends on the RNA helicase RIG-I and its paralogs. RIG-I senses double-stranded RNA (dsRNA) intermediates that are generated during the replication of single-stranded RNA (ssRNA) viruses, and initiates the production of the type 1 interferon and inflammatory cytokines via activation of the transcription factors IRF3/7 and NFKB [1,2]. Interferons can in turn activate the JAK/STAT signaling pathway to induce an antiviral state and mediate viral control. In essence, binding of interferon to the type 1 interferon receptor (IFNAR) leads to activation of the receptor associated tyrosine kinases JAK1 and TYK2, and eventually to the phosphorylation of the STAT transcription factors. Upon dimerization, STAT transcription factors can undergo translocation to the nucleus where they activate the expression of antiviral genes and inflammatory response genes [3]. On the other hand, antiviral RNAi relies on the processing by the endoribonuclease Dicer of long double stranded (ds) viral RNA molecules occurring during the viral cycle. These dsRNA molecules appear particularly during replication of the RNA genome by the virally encoded RNA-dependant-RNA polymerase. After processing by Dicer, the resulting small interfering RNAs (siRNA) are loaded in the RNA induced silencing complex (RISC) through the binding of the siRNA to a protein of the Argonaute family. Recognition of the target viral RNA by the active RISC eventually leads to its degradation [4].

In principle, all the components required for RNAi are still present in higher vertebrates, but this might simply reflect other biological functions such as microRNA (miRNA) based gene regulation. It has also been proposed that the IFN based innate immunity and antiviral RNAi might be incompatible [5,6] or that the evolution of Dicer in mammalian somatic cells renders it inactive for long dsRNA processing [7].

However, some evidence supports potential roles RNAi based antiviral immunity in mammals [8-10]. Similarly, the prominence of the RNAi pathway in fighting viruses in invertebrates may obscure other mechanisms that the innate immune system may use to combat viruses in these organisms. One such example is the nematode *Caenorhabditis elegans* (*C. elegans*), where antiviral immunity has previously been shown to involve a potent RNAi response [11].

A single virus, the Orsay virus, has been so far described to infect *C. elegans* in the wild [11]. This small bipartite single stranded RNA virus is efficiently targeted by the nematode RNAi machinery to prevent its replication. Surprisingly, *C. elegans* antiviral immunity requires DRH-1, A conserved helicase related to RIG-I, to initiate the antiviral RNAi pathway (Fig 1A) [12,13]. Interestingly, a previous analysis of gene expression changes upon viral infection revealed evidence for the induction of antiviral response genes upon viral infection independent of the RNAi pathway [14]. However, neither the upstream signaling pathway linking these gene expression changes to viral infection, nor the extent to which gene expression alterations directly contribute to antiviral defence are known. We therefore set out to uncover new signaling pathways involved sensing and transducing viral infection.

## Results

### Identification of a STAT transcription factor as a modulator of the infection response

In order to identify key factors regulating the immune state of *C. elegans* after infection by the Orsay virus (OrV), we set out to identify conserved regulatory motifs for the genes that might be involved in a transcriptional response to infection. Previously we identified a set of such putative antiviral response genes that were upregulated upon infection with the Orsay Virus in the laboratory reference strain N2 [14]. Interestingly, these genes were not upregulated in another domesticated strain of *C. elegans*, JU1580, which is hypersensitive to the Orsay Virus [12,14]. The JU1580 strain lacks the *C. elegans* RIG-I homologue DRH-1 (Fig 1A). This suggested that DRH-1 and/or signaling might act upstream of a transcriptional response to infection. However, as JU1580 and N2 differ at many loci besides *drh-1*, including some infection response genes [11,12], we decided to test this further by performing microarray analysis to compare infection induced genes in N2, a *drh-1* knockout in the N2 strain, JU1580, and JU1580 carrying a transgene containing the N2 *drh-1* locus (JU1580 *drh-1* rescue). Hierarchical clustering of differentially regulated genes recapitulated our earlier findings allowing us to identify a set of genes upregulated upon viral infection that are likely dependent on DRH-1 activity (S1 Fig). Thus, we conclude that there is a specific transcriptional response to viral infection downstream of viral RNA recognition.

**Fig 1.**
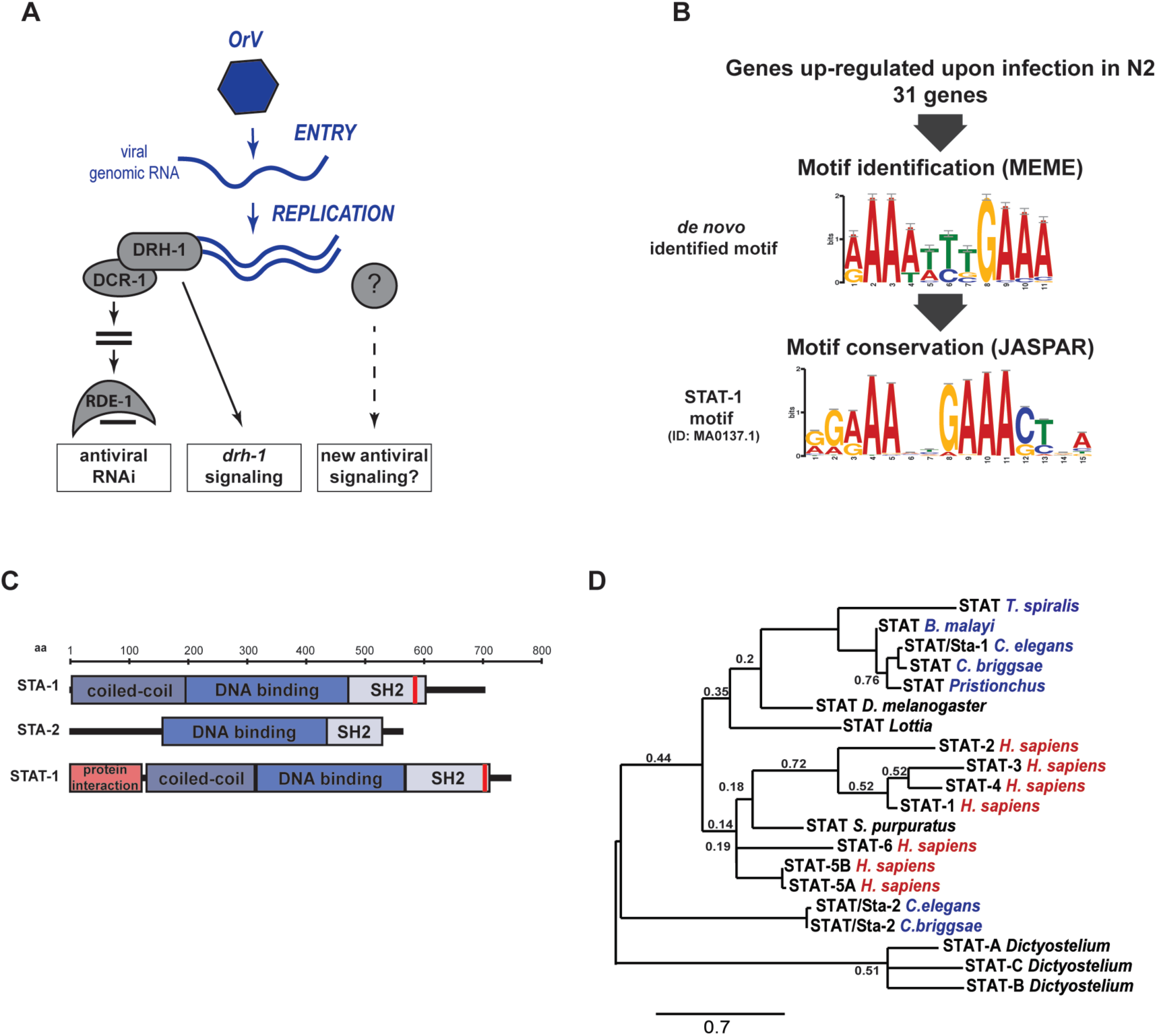
Identification of a STAT transcription factor signature in immune response against the Orsay virus in *C. elegans.* (A) Schematic representation of known antiviral pathways in *C. elegans.* (B) MEME motif enrichment in genes regulated upon infection and conservation of the *de novo* identified motif by TOMTOM. (C) Phylogenetic analysis of STAT transcription factors. Nematodes are highlighted in blue and vertebrates in red. The tree is based on full length sequences. Alignment was performed with Muscle. Branch support values are from bootstrap using 1000 iterations. Only values lower than 0.8 are depicted. (D) Schematic representation of conserved domains between STAT-1 and *C. elegans* STATs.

To get at the nature of the signaling downstream of viral recognition, DRH-1- dependent or other, we extracted the promoters from the set of genes upregulated in N2 after infection and searched for associated motifs using the motif-prediction software MEME [15]. We then compared these motifs to known transcription factor DNA binding motifs in the JASPAR core database using Tomtom [16,17]. Remarkably, we identified an enriched motif with similarity to the one of STAT transcription factors (Fig 1B), well known to have a conserved role in antiviral defence in mammals. There are two STAT homologs, STA-1 and STA-2 in *C. elegans* (Fig 1C). STA-1 has all classical functional STAT domains: a coiled-coil domain for protein-protein interaction, a DNA binding domain, a SH2 domain and a putative tyrosine phosphorylation motif [18]. STA-2 is similar but lacks the coiled-coil domain as well as the tyrosine phosphorylation motif [19]. Consistent with this, a phylogenetic tree with *C. elegans* STA-1 and STA-2 and representative STATs from other organisms suggests that STA-1 is closely related to mammalian STATs but that STA-2 is more divergent (Fig 1D). Interestingly, STA-2 has previously been implicated in an antifungal response [19], whereas STA-1 has been linked to developmental signaling in *C. elegans* [20]. We therefore speculated that STA-1 and/or STA-2 might have a previously unappreciated role in antiviral defence.

To evaluate a potential role for STAT signaling in antiviral defence in *C. elegans*, we used RT-qPCR to quantify the viral loads of *sta-1* and *sta-2* mutants following infection by the Orsay virus. As controls we used wild-type N2 animals and *rde-1* mutants, which are hypersensitive to infection due to the lack of the Argonaute protein RDE-1, which is essential for the initiation of antiviral RNAi (Fig 1A) [11,21]. Surprisingly, *sta-1* mutants were 100-fold more resistant to infection than wild-type animals, whereas *sta-2* mutants showed normal sensitivity (Fig 2A). Double mutants lacking both STAT homologs were no more sensitive than *sta-1* single mutants. We conclude that STA-1 but not STA-2 acts in regulating antiviral defense. Consistent with these observations of viral load, induction of a viral response reporter gene, comprising *gfp* driven by the promoter of the viral response gene *sdz-6* [14], was reduced in *sta-1* mutants compared to N2 and *rde-1* mutants (S2 Fig). Next we generated a single-copy, intrachromosomal *sur-5::gfp::sta-1* transgene, which drives expression of a GFP-STA-1 fusion protein in the intestine and other somatic tissues (Fig 2B). GFP-STA-1 accumulated on chromatin in the nuclei of intestinal cells (Fig 2B). Importantly, the *sur-5::gfp::sta-1* transgene restored normal sensitivity to the Orsay virus in a strain lacking endogenous *sta-1* (Fig 2C). Finally, we asked if STA-1 acts independently of the antiviral RNAi pathway using epistasis analysis (Fig 2D). In *rde-1* mutants lacking the antiviral RNAi response (Fig 1A) the Orsay virus accumulates to much higher levels than in the wild-type N2 strain. However, *sta-1; rde-1* double loss-of-function mutants show significantly reduced viral loads as compared to *rde-1* single mutants (Fig 2D) suggesting that STA-1 acts, at least in part, independently of the RNAi pathway in viral infection (Fig 2D). Together, these results identify the *C. elegans* STAT transcription factor STA-1 as acting in parallel to RNAi in the immune response to viral infection in *C. elegans.*

**Fig 2.**
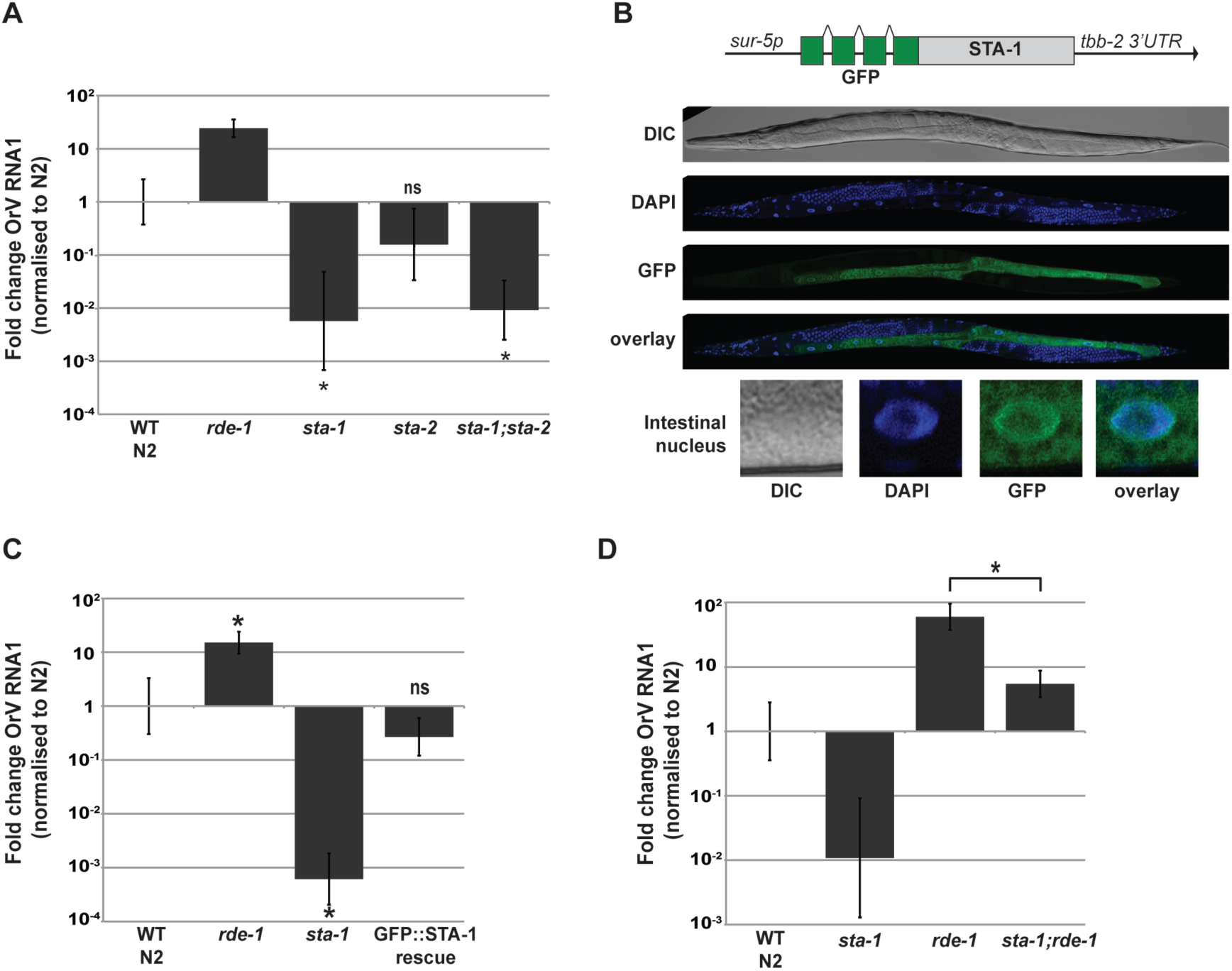
STA-1 is a key transcription factor in the immune response upon Orsay virus infection. (A), (C), (D) Viral load in different strains, measured by RT-qPCR on the Orsay virus RNA1 genome after 3 days of infection. *= p<0.01, two-tailed Mann-Whitney U test. The infection were performed on the control N2 and *rde-1* (ne219) strains, and strains with the following mutations: *sta-1* (*ok587*)*, sta-2(ok1860).* (B) Expression of a single copy transgene coding for a GFP::STA-1 fusion protein under the control of a *sur-5* promoter in adult animals.

### STA-1 is a repressor of infection response genes

In order to test whether the hyper-resistance of *sta-1* mutants to viral infection reflects a role for STA-1 in the regulation of antiviral response genes we performed RNAseq analysis of wild-type N2 and *sta-1* mutant animals with or without OrV infection (Fig 3A). We then selected transcripts altered significantly after OrV infection (DESeq, q <0.1). We observed a robust response to infection in N2 animals, whereas fewer transcripts were significantly altered in *sta-1* mutants upon infection (Fig 3B). However, *sta-1* mutants displayed a constitutive de-regulation of gene expression as compared to N2 animals in the absence of OrV infection (Fig 3B).

**Fig 3.**
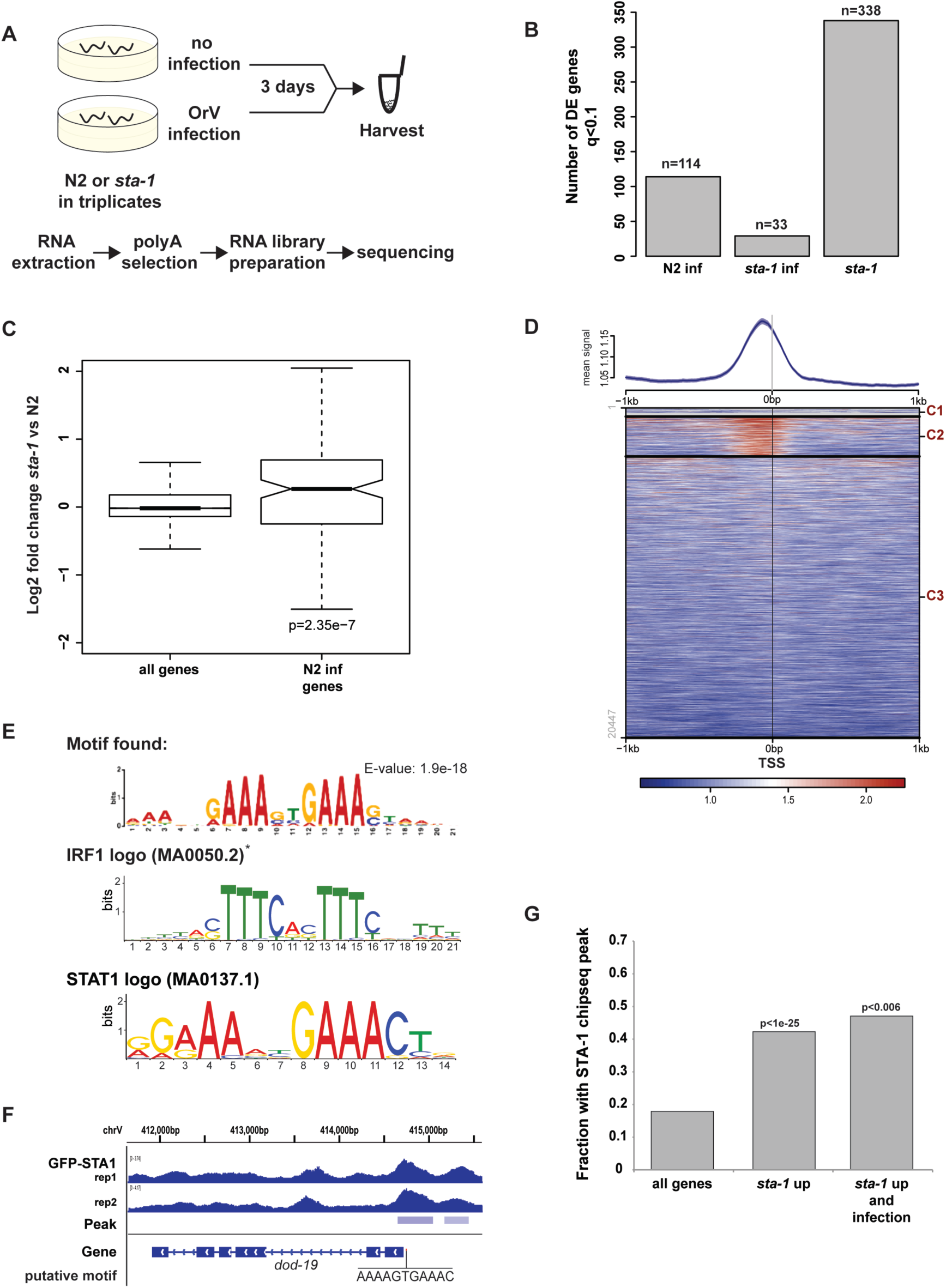
STA-1 acts at the promoter of virus response genes and represses their expression. (A) Schematic representation of the RNAseq experiment performed with N2 and sta-1(ok587) animals. (B) Representation of the number of genes showing differential expression upon infection in N2, infection in *sta-1* mutant animals or in *sta-1* mutants compared to N2, as measured by RNAseq. (C) Gene expression measured by RNA-seq in a *sta-1* mutant animals, normalized to N2 control. All genes or a subset of genes that show upregulation upon infection in N2 animals are depicted. Box shows interquartile range and the whiskers extend to the greatest value <=1.5 times the interquartile range from the box. The significance of the enrichment is represented as the p-value of the Wilcoxon unpaired test. (D) ChIPsequencing after immunoprecipitation of the GFP::STA-1 transgene. On the top part, the ChIP signal is plotted 1kb upstream and 1kb downstream of all genes anchored at TSS as in [43,44]. The heat map was generated using k-means clustering (3 clusters) of all TSS using the GFP::STA-1 signal for clustering. (E) Motif identification and conservation by Centrimo [45] for the 1 kb sequence centered around the summits of the peaks in the cluster 2 (C2) in (D). * IRF1 motif identified by Tomtom to match the reverse complement of the motif found. (F) Example of a *sta-1* upregulated gene showing binding of STA-1 by ChIP seq in two independent biological replicates. The putative STA-1 binding motif is indicated. (G) ChIP peaks enrichment in different sets of genes. Genome wide (all genes), genes upregulated in *sta-1* vs N2 *(sta-1* up) and genes differentially expressed upon infection and upregulated in *sta-1 (sta-1* up and infection).

Furthermore, genes that changed expression significantly upon infection in N2 showed a strong trend to be constitutively upregulated in *sta-1* mutants, while this was not the case when considering all genes (Fig 3C). These data suggest that STA-1 largely acts as a transcriptional repressor of an antiviral gene expression program. However, this antiviral gene expression program includes genes that are up- and downregulated upon infection, either directly or indirectly (Fig 3B). We therefore propose that the increased resistance of *sta-1* mutants to viral infection is caused by a constitutive antiviral defence genes expression program.

Next we asked whether STA-1, similarly to STATs in mammals, might bind to DNA in a sequence specific manner to regulate antiviral response genes. We therefore performed ChIPseq analysis of GFP::STA-1 to determine its genomic binding pattern in animals expressing the *sur-5::gfp::sta-1* transgene but lacking endogenous *sta-1.* Through GFP::STA-1 ChIPseq, we identified 2133 STA-1 peaks across the genome. STA-1 was enriched close to transcription start sites (TSS) of about 20% of genes with an enriched binding peak ~200bp upstream of the TSS (Fig 3D). These data are in agreement with STA-1 acting as a specific DNA-binding transcription factor to regulate gene expression. To test further the association of STA-1 binding with the antiviral gene expression response we used the sequences surrounding the STA-1 peaks in order to search for enriched motifs. Importantly, we recovered an enriched motif that was nearly identical to the motif predicted from our gene expression analysis (Fig 1B) and matching the mammalian IRF and STAT motifs in the JASPAR core database and the consensus core interferon sensitive response element (ISRE) TTCNNTTT (Fig 3E, 3F) [22]. Intriguingly, we additionally identified a separate highly enriched motif with strong similarity to the consensus sequence for GATA-like transcription factors, suggesting that a GATA protein may be a cofactor for STA-1 at some of its binding sites (S4 Fig). The predicted GATA and STAT motifs did not intersect suggesting that GATA-like transcription factors may be able to recruit STA-1 to DNA independently of STA-1 DNA binding, consistent with previous studies in mammalian cells [23].

We next tested the association between STA-1 DNA binding and gene expression. STA-1 binding was strongly enriched at genes with increased expression in *sta-1* mutant animals (p<1e-25, Fig 3G, S3 Fig), as compared to all genes. This confirmed our earlier observation that STA-1 largely acts as a constitutive repressor of gene expression. The intersection of STA-1 binding with all N2 infection response genes including those not upregulated in *sta-1* mutants was not significant; however, STA-1 binding was enriched strongly at N2 response genes that were also upregulated in *sta-1* mutants, although these were few in number (Fig 3G).

### STA-1 is required for normal lifespan

The hyper-resistance of *sta-1* mutants to infection raises the question of whether there are negative fitness consequences associated with STA-1 deficiency, that might act as trade-offs between resistance to infection and optimal growth. No obvious defects on development or fecundity were observed in *sta-1* mutants, consistent with published data [18]. However, we observed a significant decrease in the median lifespan of *sta-1* mutants (Fig 4). This may be due to the constitutive activation of pathogen response genes in *sta-1* mutants imposing a cost on animal development and/or physiology as has been shown in other systems, e.g. insects [10].

**Fig 4.**
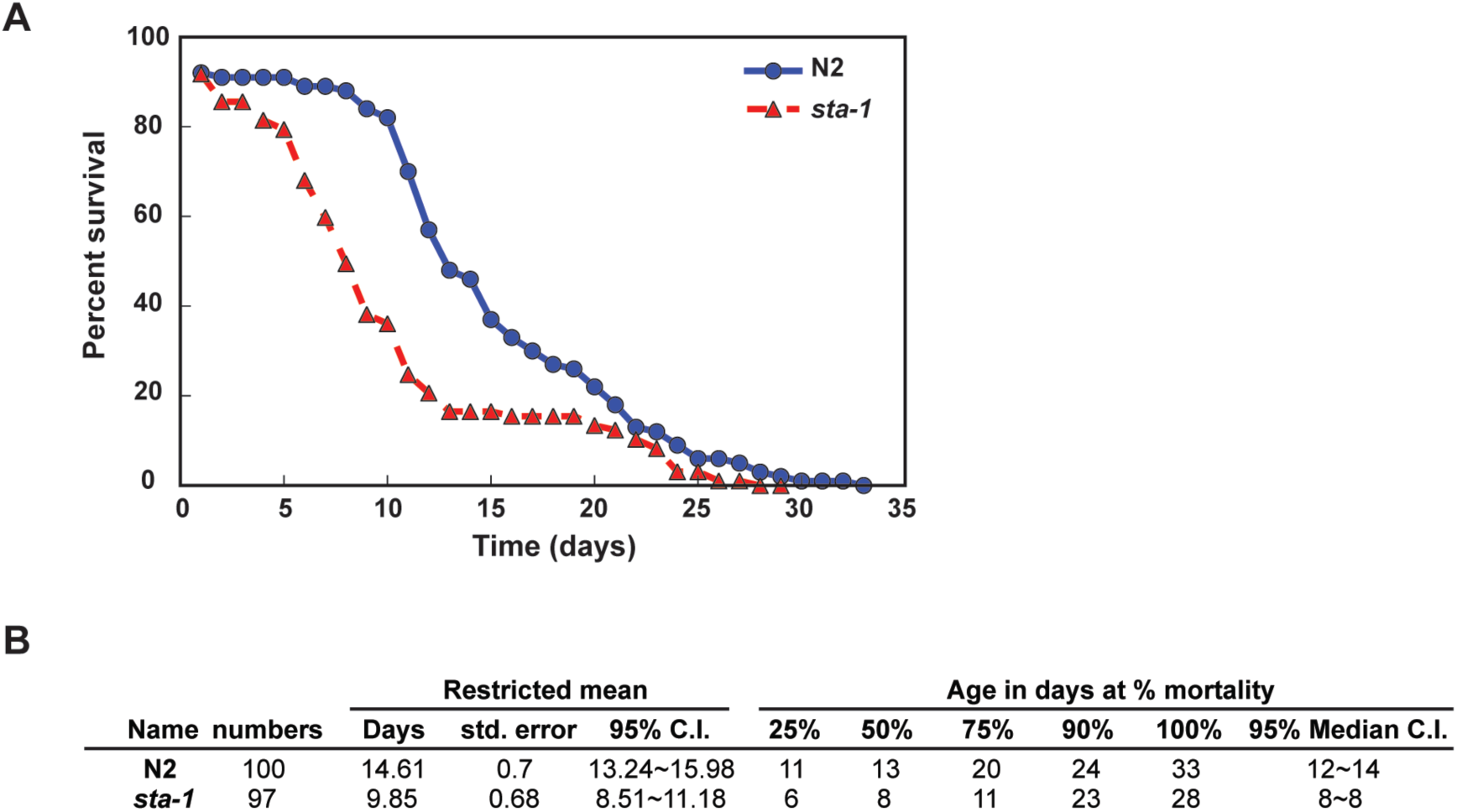
STA-1 is required for normal lifespan. (A) Survival plot showing the relationship between the survival and age of the wild-type N2 and *sta-1* mutant animals. (B) Descriptive statistics of the lifespan experiment in (A). The associated Log-Rank test Bonferroni p-value is 2.7e-06.

### The SID-3 kinase acts upstream of STA-1 in regulation of the accumulation of the Orsay virus

Having demonstrated that STA-1 acts as a constitutive repressor of antiviral response genes, we wondered how STA-1 activity was inhibited in response to viral infection. In mammals STAT proteins are regulated by JAK tyrosine kinase phosphorylation (Fig 5A). There is no conserved homolog of the JAK kinases in *C. elegans*, however the canonical tyrosine phosphorylation site on STA-1 is conserved. Thus other kinases may regulate STA-1 activity [18]. There is also a growing body of evidence of JAK-independent phosphorylation of STAT transcription factors, including serine phosphorylation [24,25]. Additionally, tyrosine and serine/threonine kinases play an essential role in antibacterial and antifungal defence in *C. elegans* [26,27]. To identify potential kinases upstream of STA-1, we performed an RNAi screen, testing genes with the serine/threonine/tyrosine-protein kinase catalytic domain IPR001245 [28]. The *C. elegans* genome encodes 176 proteins with this domain, 116 of which were available as clones as part of *C. elegans* genome-wide RNAi libraries [29,30]. Following RNAi by feeding, we infected animals with the Orsay virus and then quantified viral load for each of the 116 candidates and additional controls (Fig 5B, 5C). Using a stringent cut-off (|Z-score| > 3) we only identified a single regulator of viral load, *sid-3.* RNAi of *sid-3* resulted in ~100- fold reduction of viral RNA accumulation compared to control (Fig 5C). We confirmed that *sid-3* is required for sensitivity to viral infection by testing two independent deletion mutants of *sid-3* (Fig 6A, 6B). SID-3 is a tyrosine kinase, implicated in systemic RNAi, and is presumed to assist in the import of dsRNA into the cell during experimental RNAi [31]. However, this role in RNAi is not likely to be linked to its role in antiviral defence, because other genes required for systemic RNAi such as *sid-1, sid-2* and *sid-5* do not show a significant difference in antiviral sensitivity from wild-type N2 ([32], Fig 6C).

**Fig 5.**
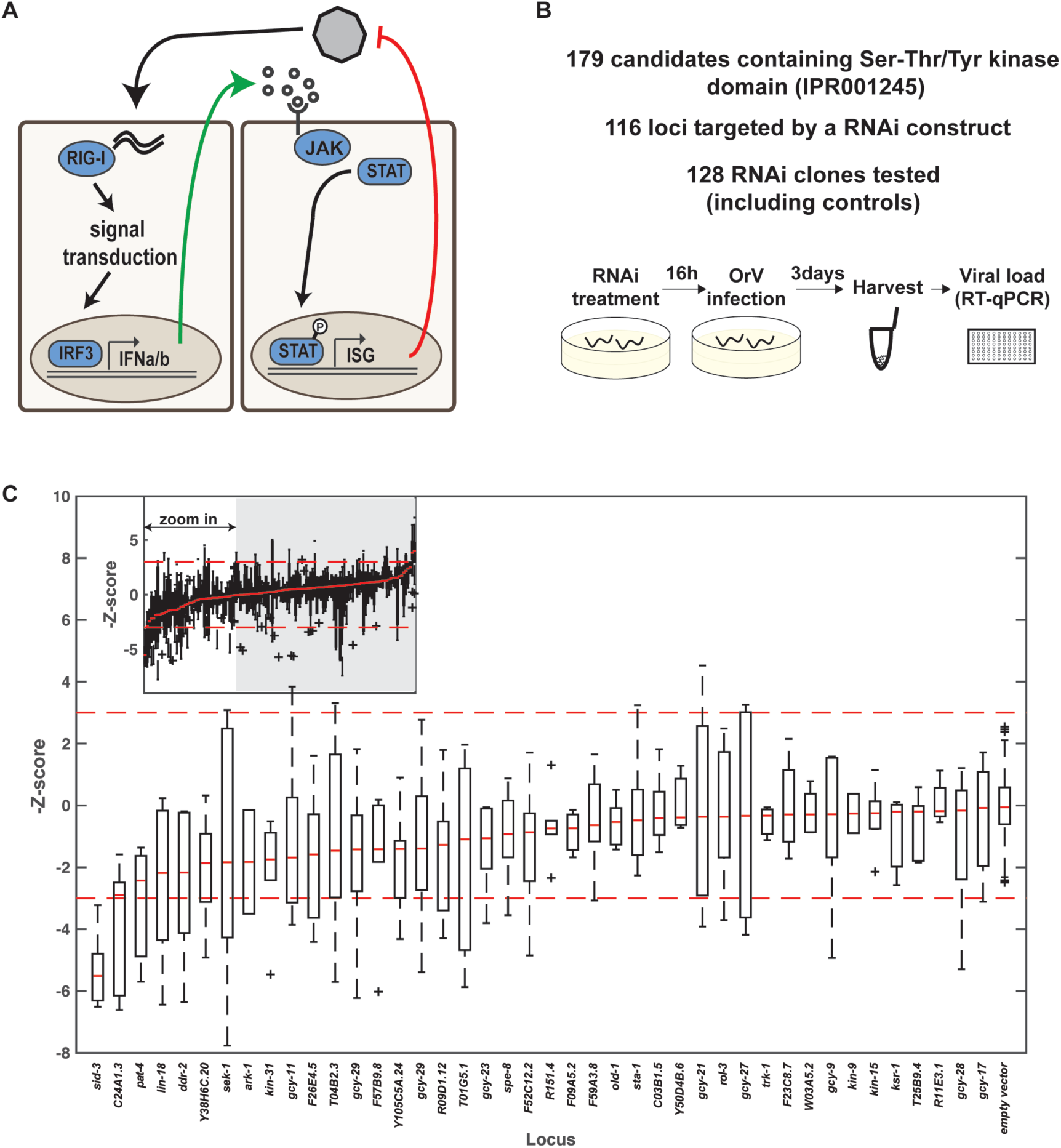
The tyrosine kinase SID-3 enables efficient viral replication. (A) Activation of the JAK-STAT signaling pathway in RNA virus infection in humans. Upon recognition of the viral RNA by sensor like RIG-I, type 1 interferon (IFN α/β) are released in the environment, recognised by the IFN receptor and activate the JAK tyrosine kinase to trigger nuclear translocation of activated STAT transcription factors. IFN: Interferon, ISG: Interferon stimulated Genes. (B) Overview of the RNAi screen. (C) The viral RNA accumulation after RNAi treatment is represented by the z-score of the ΔΔCt values (relative to empty vector, eV). The dotted lines represent the 99% confidence interval of the control RNAi (eV). The red bars represent the median of the biological replicates for the genes tested or for controls. The inlet at the top depicts all tested clones, the main figure represents RNAi treatment leading to increased resistance to OrV infection.

**Fig 6.**
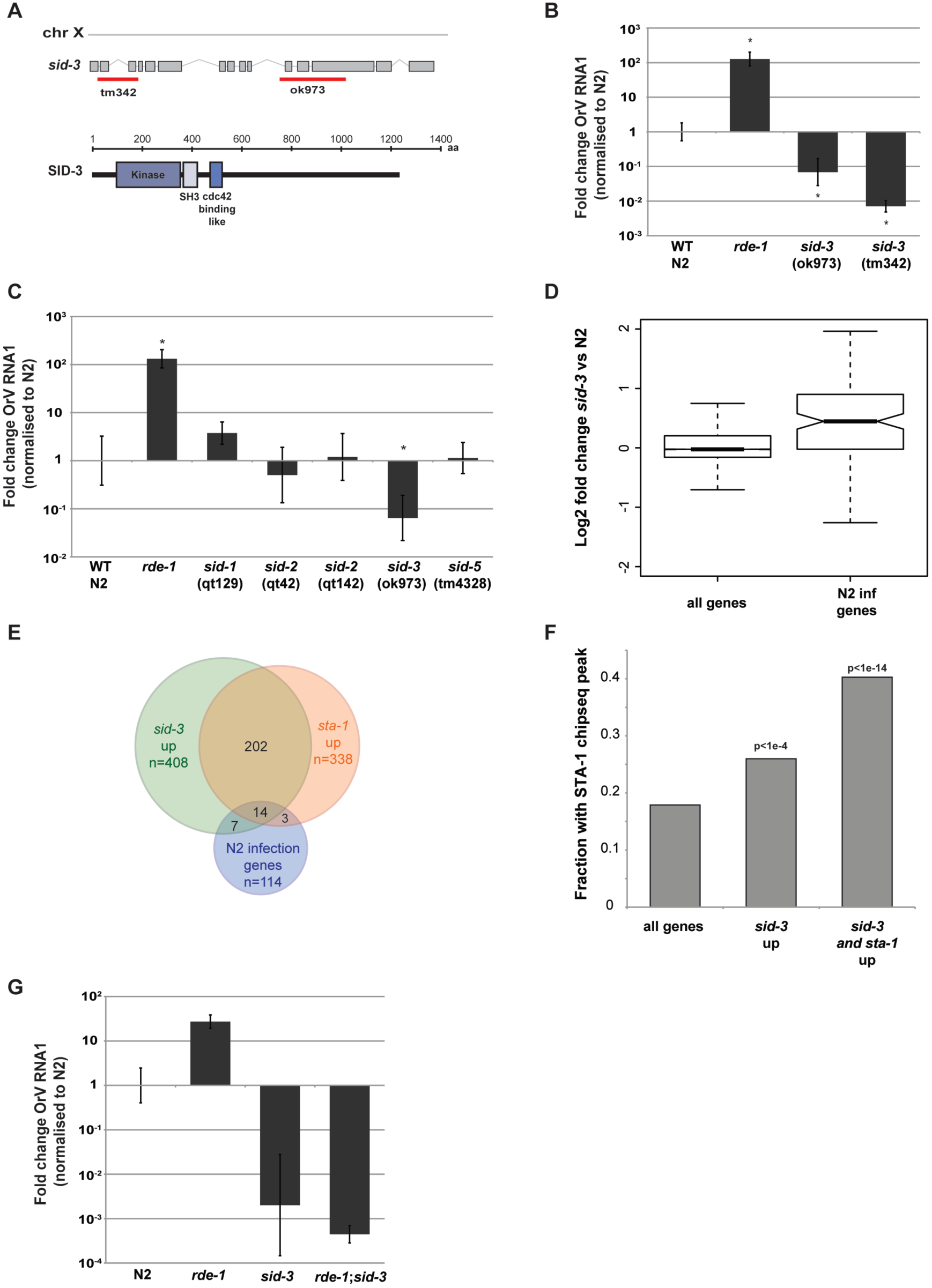
SID-3 and STA-1 regulate a common set of genes. (A) Schematic representation of the different *sid-3* alleles available. *sid-3* ok973 allele induces a deletion of 1330 bp and insertion of 12 nucleotides, deleting exons 11,12 and part of exon 13, after the tyrosine kinase domain. *sid-3* tm342 is a null allele, where an early 821 bp long deletion (exon1 to exon3) and insertion of 7 nucleotides, lead to a premature stop codon. (B), (C) Viral load after 3 days of infection, measured by RT-qPCR on the Orsay virus RNA1 genome. Mann-Whitney test, two-tailed, *= p<0.01. (D) RNA-seq analysis showing an enrichment for infection genes in *sid-3*(ok973) upregulated genes. (E) Overlap between infection genes and *sta-1* or *sid-3* upregulated genes. (F) Overlap between STA-1 bound genes, as shown by ChIP-seq and *sid-3* upregulated genes. (G) Viral load in the indicated strains, measured by RT-qPCR on the Orsay virus RNA1 genome after 3 days of infection. The *sid-3* allele used is *sid-3*(ok973).

We therefore wondered whether SID-3 might act in the same pathway as STA-1. We quantified gene expression in *sid-3* mutants using RNAseq. Similar to what we observed for *sta-1* mutants, genes that changed expression significantly upon infection in N2 showed a strong trend to be constitutively upregulated in *sid-3* mutants, while this was not the case when considering all genes (Fig 6D). Furthermore, the genes upregulated in *sid-3* mutant animals showed a striking overlap with the genes upregulated in *sta-1* mutant animals (p=1.2e-15). Additionally, *sid-3* upregulated genes were enriched for antiviral response genes (p=5.55e-10) (Fig 6E and S3 Fig). Moreover, genes upregulated in *sid-3* mutants, including those shared with *sta-1*, were enriched for STA-1 binding by ChIPseq, suggesting that *sid-3* acts upstream of *sta-1* in the antiviral gene expression response (Fig 6F). We conclude that SID-3 and STA-1 act in the same pathway to regulate an innate antiviral immunity program.

Finally, we addressed the relationship between SID-3 and the antiviral RNAi pathway using epistasis analysis. Interestingly, *sid-3; rde-1* double loss-of-function mutants were as resistant to OrV infection as *sid-3* mutants (Fig 6G). This is in contrast to what we observed in *sta-1;rde-1* mutants (Fig 3D). This suggests that SID-3 might act at an extremely early stage of viral infection.

We conclude that SID-3 acts both upstream of antiviral RNAi and upstream of a STA-1-dependent antiviral gene expression program to regulate the response to OrV infection. Together, these data are in support of a model (Fig 7) whereby in uninfected animals, SID-3 signals to STA-1, potentially by phosphorylation, to maintain repression of antiviral response genes. Upon infection, this signaling is curtailed, leading to loss of repression and upregulation of antiviral response genes.

**Fig 7.**
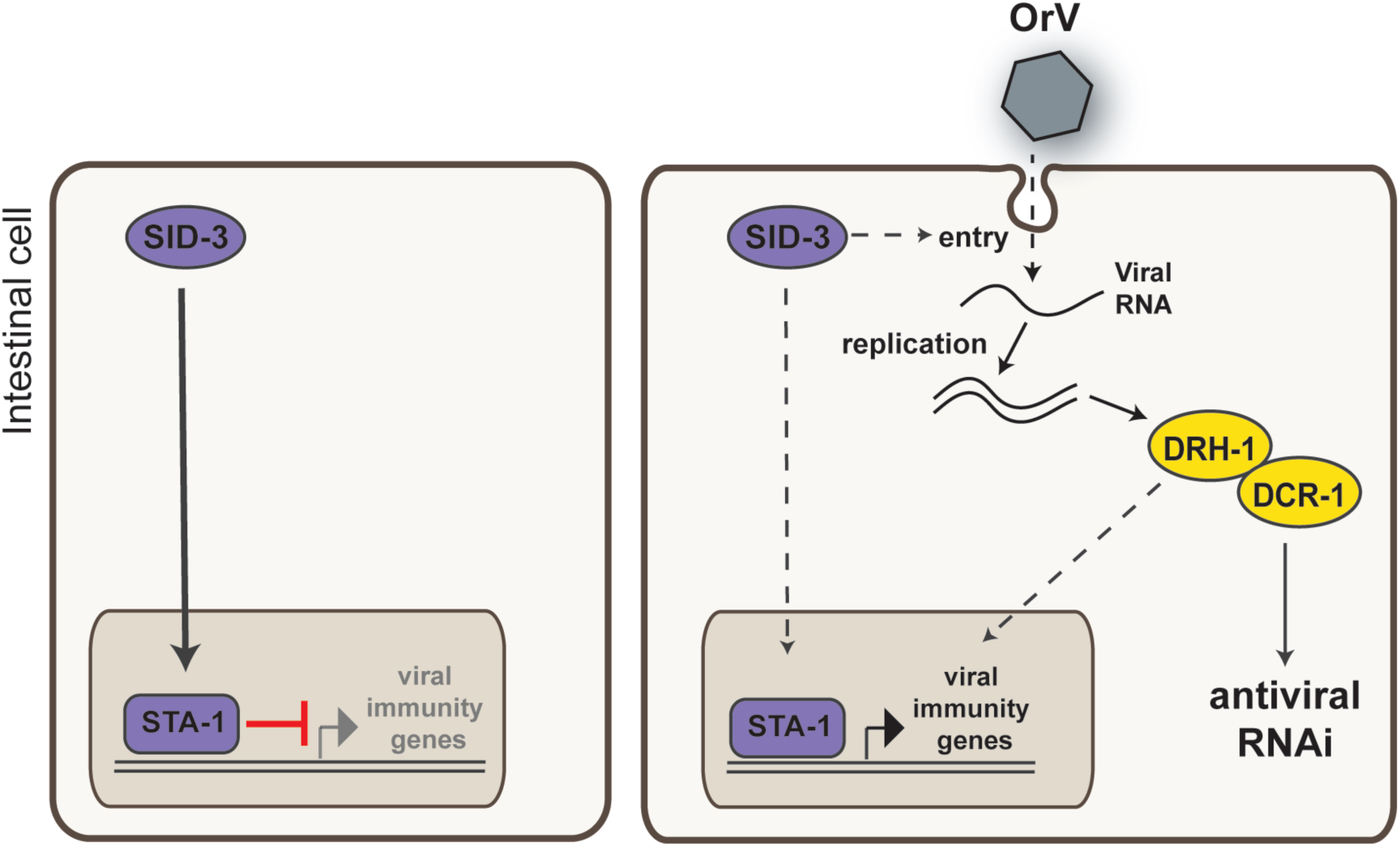
A STA-1 pathway controls the response to viral infection in *C. elegans*, a model.

## Discussion

Here, we uncover a hitherto unappreciated role for the *C. elegans* homologue of the mammalian STAT family of transcription factors. We demonstrate that the tyrosine kinase SID-3 and the STAT transcription factor STA-1 are tightly linked functionally. Additionally, we uncover a novel potential signaling pathway linking STAT activity to viral RNA recognition. Our results have intriguing implications both for innate immunity in *C. elegans* and for the evolution of eukaryotic signaling pathways.

### Regulation of antiviral gene expression in *C. elegans*

Previously there has been some debate over the extent to which gene expression responses to infection in *C. elegans* represent specific pathogen response pathways or more general responses to stress. Our finding that a STAT-family transcription factor is responsible for constitutively repressing antiviral response genes, and that this is relieved upon infection, suggests that at least some part of the antiviral gene response is a specific response. Moreover, constitutive upregulation of these genes, as demonstrated by mutants lacking *sta-1*, leads to hyper-resistance to viral infection indicating that at least some of these antiviral response genes play direct roles in protecting *C. elegans* from viral infection. Further examination of the functions of these genes may lead to the identification of novel mechanisms underlying their antiviral activity. How is the constitutive repression of antiviral response genes by *sta-1* relieved upon viral infection? The fact that *sid-3* mutants show loss of *sta-1* repression suggests that phosphorylation of STA-1 downstream of SID-3 is required for it to exert its repressive function. Thus, viral infection may lead to loss of SID-3 activity and concomitant loss of STA-1 phosphorylation. Exactly how SID-3 activity is downregulated upon viral infection is unclear. However, an interesting clue to the potential role of SID-3 in viral infection comes from its proposed role in endocytosis [31]. Non-enveloped viruses require the endosomal pathway for entry into cells; it is therefore plausible viral entry into endosomes may lead to titration of SID-3 away from its role in signaling to STA-1, leading to relief of repression. Such an early role for *sid-3* in infection is consistent with the ability of *sid-3* mutation to suppress the sensitivity of RNAi pathway mutants. Indeed, in a co-submitted paper, Dave Wang and colleagues demonstrate that *sid-3* is required for viral entry. Thus, we hypothesise that the dependence of the virus on *sid-3* for its entry has been exploited by *C. elegans* as a mechanism to regulate antiviral gene expression. Further work, potentially testing the role of other kinases we identified in our screen, will be required to complete this signaling pathway.

### Evolution of STAT signaling

It is remarkable that the STAT-family of transcription factors has a role in antiviral gene regulation in both mammals and *C. elegans*, despite the fact that both upstream and downstream genes in the pathway are not conserved between the two. Even more remarkably, the mechanism whereby STAT transcription factors contribute to gene expression regulation are quite distinct in nematodes from mammals, as whilst STA-1 is constitutively bound to DNA and acts as a transcriptional repressor in *C. elegans*, STATs are activated in response to infection. Despite this, there appear to be intriguing parallels between STA-1 activity and the mammalian STAT pathway, in particular the association that we uncovered between STA-1 binding and GATA-transcription and the conserved role of STATs in viral gene expression responses in *C. elegans* and mammals.

Our results highlight the remarkable plasticity of signaling pathways through evolution, whereby a central module may retain the same function over millions of years whilst acquiring distinct regulatory partners. This ability to both retain and diversify functionality simultaneously is clearly particularly important in the immune system, where the specific nature of the threats involved changes rapidly whilst their general modes of action may not. It will be intriguing to explore whether STATs in other metazoans have similarly diversified whilst retaining antiviral activity- it is possible that the *C. elegans* mode of STAT action that we uncover here is in fact the original conserved role of STAT transcription factors in viral responses. If so, vestiges of this function may be retained in mammalian cells, particularly those without expression of the JAK pathway.

## Materials and Methods

### Datasets

High-throughput sequencing datasets are accessible through the GEO repository.

### Nematode culture and strains

We grew *C. elegans* under standard conditions at 20°C. The wild type strain was var. Bristol N2. The food source used was *E. coli* strain HB101 (*Caenorhabditis* Genetics Center, University of Minnesota, Twin Cities, MN, USA). Detailed information about all strains generated and used in this study are in the S1 Table.

### Viral filtrate

Stably infected populations of sensitive animals (JU1580) were transferred to 2 L liquid culture with HB101 food and grown for 7 days. The supernatant of the culture was harvested on ice and filtered with a 0.22 μm filter. The resulting viral filtrate was aliquoted and stored at −80°c.

### Viral infection

Two WT or three mutants L4 animals were added on seeded 50 mm plates. 16 hours later, 20 μl of Orsay virus filtrate was added to the edge of the bacterial lawn. Animals were collected 3 days after infection. The animals were washed off the plates with M9 buffer. The animals pellets were washed another three times by pelleting the animals either by gravity on ice or by centrifugation at 800 g for 2 minutes in a swinging bucket centrifuge, snap-freeze in liquid nitrogen and stored at −80°C.

### Lifespan assay

100 animals of each strain were picked onto ten 50mm NGM plates. Adults animals were transferred every 2 days onto fresh plates for the duration of egg laying. Their survival was measured by movement of the head. Animals showing no head movement were gently touched twice with a worm pick and observed for 30 seconds following each touch. Animals not moving were considered dead and removed from the plate. The survival curves were plotted and analysed using OASIS (Online Application for the Survival Analysis) [33].

The experiment was reproduced 3 times with one representative example illustrated.

### RT-qPCR

The worm pellets lysis was performed with 5 μl of worm pellet and 45 μl of the lysis solution from the Power SYBR Green Cells-to-Ct kit (Ambion). Ten freeze-thaw cycles and 30 minutes shaking at room temperature were performed before the lysis incubation step. The RT-qPCR was performed according to the manufacturer’s instructions. The qPCR was run on a Step One Plus Real Time PCR system (Applied Biosystems). The analysis was done using the ΔΔCt method. The Ct values of 4 to 6 biological replicates per experiments (as indicated) were pooled to generate the average ΔCt for each strain and an associated standard error (sd). The ΔΔCt value was calculated with N2ΔCt as the calibrator. The 2^^^(-ΔΔCt) was plotted as the fold change and the interval of confidence is represented by error bars as 2^(- (ΔΔCt+sd(ΔCt))) and 2^(-(ΔΔCt-sd(ΔCt))). The significance of the differences observed were tested by a two-tailed Mann-Whitney *U* test at a significance level of p<0.01.

### DNA constructs

The viral sensor construct and the *gfp::sta-1* construct were generated by Gateway cloning using Multisite Gateway Three-Fragment vector construction kit (Life Technologies). Gateway entry clones containing each of the following were generated by standard techniques: *sur-5* promoter, *sdz-6* promoter, *sta-1* coding sequence, eGFP(F64L/S65T), *tbb-2* 3’UTR. Details on cloning and plasmid sequences are available upon request. The single-copy transgene was generated by transposase-mediated integration (MosSCI), as described [34,35], at insertion site ttTi5605 on chromosome II. Injection mixes contained: 20 ng/μl of vector, 20 ng/μl of Mos1 transposase (only for MosSCI), and 5 ng/μl of a pharynx marker. Integration of the extrachromosomal array was performed by EMS treatment (50 mM EMS for 4 hours).

### Sensor scoring

Scoring of the GFP was made under a Leica MZ16F fluorescence stereomicroscope.

### Imaging

Adults animals were harvested from a plate, washed off in M9. After 3 additional washes in M9 buffer, the pelleted animals were fixed in 0.5 ml of pre-cooled methanol at −20°C for 10 minutes. The pellet was washed 3 times in TBS / 0.1% Tween20 (TBS-T) and then incubated for 10 minutes in a DAPI solution (0.5 mg/ml in TBS-T). The pellet of animals was washed 3 times in PBS-T. 5μL of the pelleted animals were pipetted directly onto a Cel-line diagnostic microscope slide (Thermo Scientific) and imaged using a Leica SP8 upright microscope.

### Microarray

Microarray data were processed using rma and annotations were obtained through the Bioconductor pipeline in the R programming environment. Differentially regulated genes in any condition identified using t-test, p<0.01 and a 2-fold difference upon infection. Data are available in the S1 File. Hierarchical clustering on the *drh-1* array was carried out on the set of differentially regulated genes using the hclust function in R and Ward’s method.

### Motif analysis

To specifically identify potential viral response genes we normalized the array by *cul-6* as this removed variation in the final level of infection. Qualitatively this identified similar trends to those previously published [14], such as putative antibacterial response genes upregulated specifically in JU1580 and *drh-1* upon infection. We identified genes that were >40% induced relative to *cul-6* and used BiomaRt to download the upstream sequences (S2 File).

These were used as an input to Meme attempting to find 0 or 1 site per gene. We screened these manually to avoid spurious hits and then compared potential motifs identified to the JASPAR core database. To test enrichment of STAT motifs in STA-1 ChIP-seq peaks we used FIMO to search for the STAT-1 core motif within the 500bp sequences upstream of genes containing a STA-1 binding site, applying a false-discovery rate cutoff of 0.2, which were expressed at increased level (DESeq q<0.1) in sta-1 mutants relative to N2. We compared this to a random set of genes chosen from the RNAseq data (see below).

### RNA seq

Animals were grown on 50mm NGM plates, infected or not and harvested as described in the VIral infection methods section. Total RNA was extracted using Trisure (Bioline, UK). RNA library preparation was performed with the NEBNext^®^ Ultra™ RNA Library Prep Kit for Illumina^®^ with total RNA purified with the NEBNext Poly(A) mRNA Magnetic Isolation Module, according to manufacturer’s instructions (NEB, USA).

### RNA-seq analysis

RNAseq data was aligned to the *C. elegans* transcriptome WS190 using bowtie2. Counts were obtained from resulting bam files using bedtools [36] and these were used to generate normalized data tables using DESeq [37] (S2, S3, S6 Tables). Significance between intersecting datasets was calculated by a Fisher’s exact Test (S3 Fig and S5 Fig). A cutoff of mean 25 normalised reads (normalised according to DESeq’s negative binomial distribution) for at least one condition was used and significantly altered genes were selected (DESeq Benjamini-Hochberg multiple test correction q<0.1).

### ChIP sequencing

Animals were grown on NGM plates seeded with thick HB101 food and harvested as a mixed-stage population. Frozen animals were ground to a fine powder and fixed in 1% formaldehyde/PBS for 10 minutes, quenched with 0.125 μM glycine, and then washed 3X in PBS with protease inhibitors. The pellet was resuspended in 1 ml of FA buffer (50mM HEPES/KOH pH7.5, 1 mM EDTA, 1% Triton X-100, 0.1% sodium deoxycholate, 150 mM NaCl with protease inhibitors) per 4 ml of ground worm powder. Extract was sonicated to a size range of 200-1000bp using a Diagenode Bioruptor Pico with a setting of 18 pulses, each lasting 30 seconds followed by a 30 seconds pause. The extract was spun for 10 minutes at 16000g at 4°C, and the soluble fraction was flash frozen in liquid nitrogen and stored at −80°C until use. Each ChIP was prepared in 500 μl of FA buffer containing 1% sarkosyl. 15 μg of an anti-GFP antibody (ab290) was incubated with 3 mg of extract. Additionally 10% of extract was saved as a reference. After overnight rotation at 4°C, 40 μl of blocked and washed magnetic protein A dynabeads (Invitrogen) were added and the incubation continued for 2 additional hours. Beads were washed at room temperature twice for 5 minutes in FA buffer, once in FA with 500 mM NaCl for 10 minutes, once in FA with 1 M NaCl for 5 minutes, once in TEL buffer (0.25 M LiCl, 1% NP-40, 1% sodium deoxycholate, 1 mM EDTA, 10 mM Tris-HCl, pH 8.0) 10 minutes and twice in TE pH 8.0 for 5 minutes. DNA was eluted twice with 57 μl elution buffer (1% SDS in TE with 250 mM NaCl) at 65°C, 15 minutes each time. Eluted DNA was incubated with 20 μg of RNase for 30 minutes at 37°C and then with 20 μg of Proteinase K for 1 hour at 55°C. Input DNA was also diluted in 114 μl elution buffer and treated with ChIP samples. Crosslinks were reversed overnight at 65°C. DNA was purified on PureLink PCR purification columns (Invitrogen). The libraries were prepared using a modified TruSeq ChIP Library preparation kit protocol (https://ethanomics.files.wordpress.com/2012/09/chip_truseq.pdf). The size selection was performed using Agencourt AMPure XP beads (Beckman Coulter).

### ChIP-seq analysis

*Alignment to reference genome.* Chip-seq and RNA-seq libraries were sequenced using Illumina HiSeq. Reads were aligned to the WS220/ce10 assembly of the *C. elegans* genome using BWA v. 0.7.7 [38] with default settings (BWA-backtrack algorithm). The SAMtools v. 0.1.19 ‘view’ utility was used to convert the alignments to BAM format. Normalized ChIP-seq coverage tracks were generated using the R implementation of BEADS algorithm [39,40]. *Summed ChIP-seq input* We generated summed input BAM files by combining good quality ChIP-seq input experiments from different extracts (8 experiments). The same summed inputs were used for BEADS normalisation and peak calls. *Peak calls* Initial ChIP-seq peaks were called using MACS v. 2.1.1 [41] with permissive 0.7 q-value cutoff and fragment size of 150 bp against summed ChIP-seq input. To generate combined peak calls, we used the modified IDR procedure (https://www.encodeproiect.org/software/idr/) with an IDR threshold of 0.05 to combine replicates (S4 Table). The pipeline for generating IDR peaks is avilable here: https://github.com/Przemol/biokludge/blob/master/macs2_idr/macs2_idr.ipy. *Mean signal distribution plots and heatmaps* The summarized signal profile and heatmap for GFP::STA-1 were generated using SeqPlots exploratory analyses and plotting software [42].

### RNAi screen

RNAi clones from the Ahringer library [29,30] were isolated on agar plates containing carbenicillin (50 μg/ml). Single colonies were picked in 2 ml LB + ampicillin (50 μg/ml) and grown 9 hours with shaking at 37°C. Bacteria were then seeded onto 50 mm NGM plates containing IPTG (1 mM), carbenicillin (25 μg/ml) and Fungizone (2.5 μg/ml). Two days after seeding, two L4 were added and then grown at 20°C. After 16 hours, plates were inoculated with 20 μl Orsay virus filtrate. Animals were collected after 3 days and viral relative genome copy number was measured by RT-qPCR. For each round of the screen, we used the following internal controls: N2 strain grown on empty vector (L4440 *E. coli* strain) was considered as normalization control; *drh-1* mutant fed on GFP RNAi as positive control for infection; and N2 fed on *drh-1* RNAi clone as positive control for RNAi. The full list of the RNAi clones and number of replicates used in this study is available in the S5 Table. Each replicate of the screen was performed with 3 biological replicates per RNAi clone treatment and the screen was repeated at least twice. The plates with accidental fungal contamination as well as the plates where the RNAi treatment led to embryonic lethality were removed from the analysis. The resulting exact number of biological replicates is indicated in the S5 Table. Empty vector treatment was included on each qPCR plate analysed and used as the calibrator. The ΔΔCt values were transformed as z-score, calculated as follow: zscore_i_ = (ΔΔCt_i_-μΔΔCt_emptyvector, n=161_)/stdev(ΔΔCtemptyvector). Boxplots of the Z-score for each treatment are represented in Fig 5. Individual values used for the analysis are available in the S5 Table.

## Acknowledgments

We thank David Jordan for scientific discussion and help with data visualization, Jérémie Le Pen for scientific discussion and contribution to reagents, Alyson Ashe for contribution to reagents, Alex Appert for technical support with ChIP sequencing experiment, Kay Harnish and Sylviane Moss for high-throughput sequencing. We thank the *Caenorhabditis* Genetics Center, University of Minnesota for providing some strains used in this study.

